# Intra-and Interhemispheric Signatures of Criticality at the Onset of Synchronization

**DOI:** 10.64898/2025.12.11.693654

**Authors:** Leonardo Dalla Porta, Pietro Bozzo, Marco N. Pompili, Damien Depannemaecker, Antonio J. Fontenele, Tomoki Fukai, Pierpaolo Sorrentino, Giovanni Rabuffo

## Abstract

The cerebral cortex must flexibly alternate between locally segregated activity that supports specialization and long-range interactions that enable integration. How cortical networks balance these competing demands remains unclear. We propose that fluctuations around a critical point between ordered and disordered phases provide a natural framework to understand coordinated neuronal activity across scales. Using simultaneous spiking recordings from the left and right prefrontal cortex (PFC) of freely behaving rats, we found that signatures of criticality emerged at the onset of neuronal synchronization, both locally within individual hemispheres and globally across the combined population. Over time, cortical activity explored a continuum of intra-and inter-hemispheric synchrony levels, including states in which neurons were locally desynchronized yet maintained interhemispheric coupling. A computational model operating near its critical regime reproduced these empirical patterns, capturing the characteristic relationship between local and long-range coordination. These results suggest that cortical networks achieve flexible transitions between local and global computation by fluctuating around a critical regime.

## Introduction

Cortical organization supports both specialized local processing and coordinated communication across distant regions [1– 3]. However, how the brain flexibly balances these competing demands remains a central question in systems neuroscience. In this context, brain criticality offers a theoretical framework to understand how such integration/segregation balance might be achieved in cortical networks [4]. Specifically, this framework proposes that neuronal populations operate near a critical point between order and disorder, where neuronal avalanches are observed across orders of magnitude [5], from local circuits to large-scale activations [6]. These characteristic dynamics maximize information processing [7], and can potentially explain how the brain coordinates local and global communication. Although evidence for brain criticality, at the neuronal scale, has come largely from single-area recordings [8], far less is known about how critical dynamics extend across multiple, interacting brain regions. This gap limits our understanding of whether criticality is a local property of neural circuits, a global feature of distributed networks, or yet a multiscale phenomenon that emerges from their interaction.

Recent work on high-density single-unit recordings revealed that signatures of criticality are not universal across the brain: they may depend on the neuronal subset examined [9, 10], the dimensions of activity space analyzed [11, 12], or the timescale of observations [13, 14]. Collectively, these findings suggest that criticality is not a static regime of brain activity, but a dynamic, context-dependent, and potentially region-specific feature. Key open questions remain: how do critical dynamics unfold within and between widely separated populations? Do spatially distant areas approach the critical point simultaneously, or can they tune independently? Is criticality preserved when distant populations are considered as a single global system rather than as isolated local circuits?

To address these questions, we recorded spiking activity simultaneously from the left and right prefrontal cortex (PFC) of freely behaving rats (Fig. 1A). This bilateral approach enabled us to track local dynamics within each hemisphere while also assessing the dynamics between them. By measuring population synchrony, neuronal avalanches, and interhemispheric correlations, we aimed to determine whether criticality arose independently in each region, emerged only when populations were considered together, or depended on interactions between them. This approach provided a direct way to link local fluctuations in synchrony with large-scale coordination and to test how long-range coupling shaped the expression of critical dynamics across the cortex. Finally, we used a mesoscale computational model with a known critical point to test whether the multiscale coordination observed experimentally is an intrinsic feature of networks fluctuating near criticality.

**FIG. 1.**
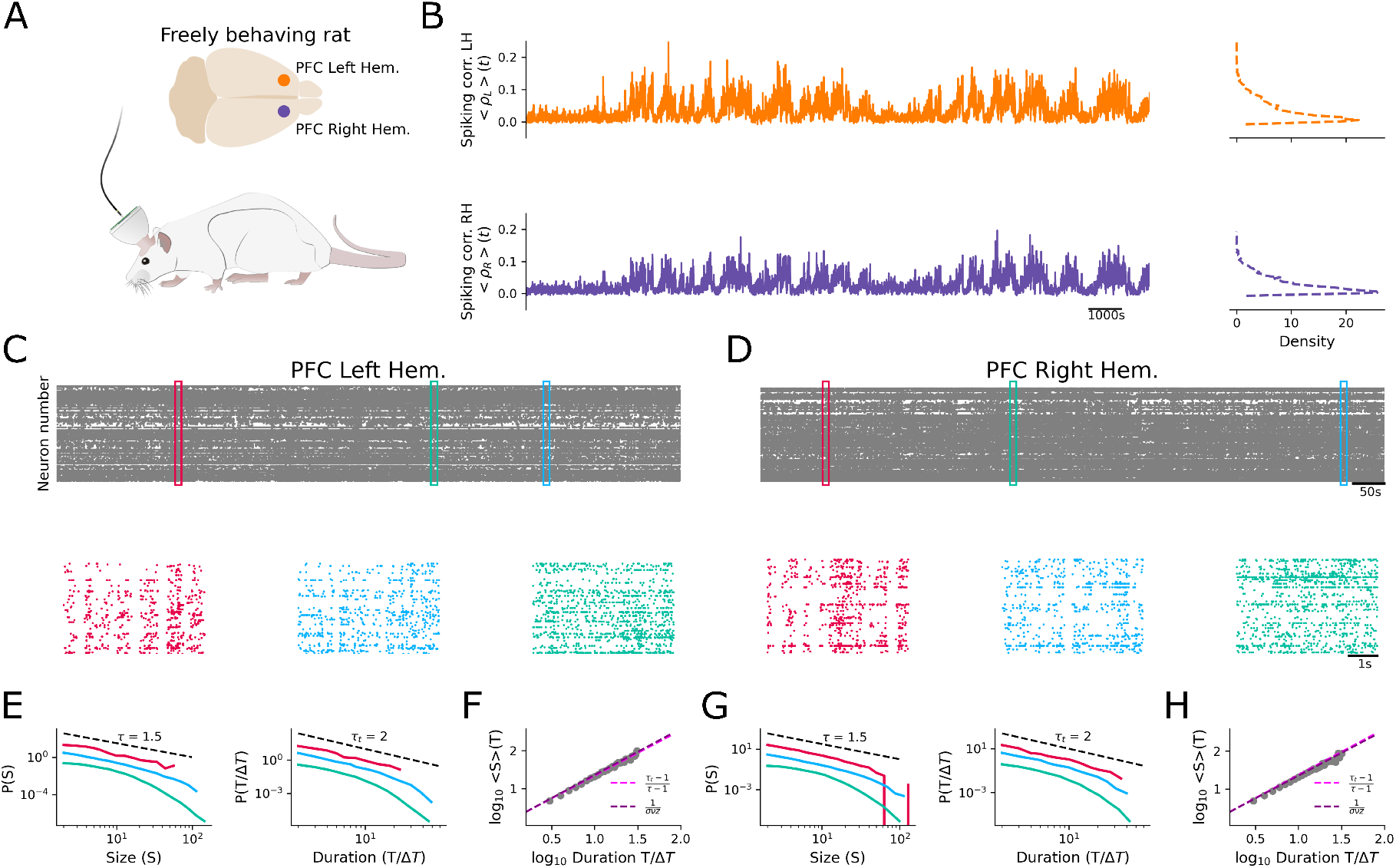
Signatures of criticality at the onset of synchronization in the freely behaving rat brain. (A) Schematics of the experiment: spiking recordings were performed simultaneously in both hemispheres of the prefrontal cortex (PFC) in freely behaving rats. (B) (Left) Mean intrahemispheric pairwise spiking correlation for left (*<ρ*_*L*_*>*(*t*)) and right (*<ρ*_*R*_*>*(*t*)) PFC regions, computed over 5-s non-overlapping time windows. (Right) *<ρ>*(*t*) histograms. (C) (Top) Raster plot of the spiking activity throughout the session. (Bottom) Zoom-in raster plot illustrating the spiking activity dynamics for different levels of synchrony: high (*<ρ>* ≃ 0.18; red), intermediate (*<ρ>* ≃ 0.09; blue), and low (*<ρ>* ≃ 0.02; green). (D) Same as in (C) but for the right PFC region. (E), (G) Neuronal avalanche distributions for size (*P*(*S*) ∼ *S*^−*τ*^) and duration 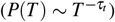 in the left (E) and right (G) PFC regions. The exponents obtained for the left (right) PFC regions during intermediate levels of synchrony were: *τ* = 1.72 (*τ* = 1.65) and *τ*_*t*_ = 1.85 (*τ*_*t*_ = 1.78). Power-law fit and exponent estimation were performed by using the methodology described in [15], setting a lower and upper bound for size (*x*_*min*_ = 3, *x*_*max*_ = 38) and duration (*x*_*min*_ = 1, *x*_*max*_ = 18). Dashed lines represent the original exponents reported in [5], which coincide with those of the mean-field directed percolation universality class [16]. The colors correspond to the states described in (C). (F), (H) Neuronal avalanches for size and duration at intermediate levels of synchrony follow the scaling relation predicted at criticality 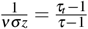 [17]. The first term (pink dashed line) is fitted from the empirical data where 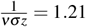 and 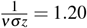 for the left (F) and right (H) regions, respectively. The second term (purple dashed line) is predicted by the relationship between avalanche exponents obtained in (E) and (G). Data shown is from one representative rat session.

## Results

### Critical Neuronal Avalanches at the Onset of Synchronization

We simultaneously recorded spiking activity from hundreds of neurons from the left and right prefrontal cortex (PFC) in freely behaving rats (n= 3; Fig. 1A). Each recording session lasted approximately 8 hours (n_*s*_=2 sessions per rat), yielding a total of 620 neurons (single unit activity, SUA): 334 neurons in the left PFC and 286 neurons in the right PFC. To quantify the level of synchrony, we computed the mean pairwise spiking correlation, *<ρ>*, in non-overlapping 5-s windows, following the same approach as in [18]. Here, the underlying assumption is that neuronal dynamics do not *rigidly* operate at a critical point; rather, they hover around a critical point, fluctuating between phases of order and disorder [18– 22].

By evaluating cortical dynamics independently in the left and right PFC populations, we observed that local population activity spontaneously alternated between periods of high and low synchrony, reflecting more ordered and more disordered states, respectively (Fig. 1B). High-synchrony periods were characterized by synchronous bursts of neuronal activations (Fig. 1C-D, red), whereas low-synchrony periods exhibited irregular firing patterns (Fig. 1C-D, green). Intermediate levels of synchrony were also observed (Fig. 1C-D, blue), during which the dynamics displayed complex spatiotemporal activity patterns known as neuronal avalanches [5]. Following the original definition [5] and the methodology of [18], we defined the size of an avalanche as the sum of spiking activity in consecutive occupied time bins (Δ*T*), with Δ*T* set to the average interspike interval [23]. Avalanche duration was determined by the number of consecutive active time bins. Avalanches were computed separately within each 5-s window and grouped according to the population synchrony level (*<ρ>*) using 30 non-overlapping bins spanning the full observed range of *<ρ>*.

Analyzing neuronal avalanches independently for each PFC population revealed that, at intermediate levels of synchrony, both the size (*P*(*S*) ∼ *S*^−*τ*^) and duration 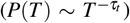 of events were well described by a power-law distribution, a hallmark signature of critical dynamics [5, 24] (Fig. 1E-G). Moreover, the scaling relation 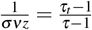, expected to hold at criticality, was also satisfied [17] (Fig. 1F-H). Here, the first term was obtained empirically by fitting 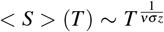 to the neuronal avalanches (Fig. 1F-H), while the second term was estimated from the fitted avalanche exponents *τ* and *τ*_*t*_ [17, 25]. *This scaling relation has been used in the literature as a stringent test of criticality [17–19], with the absolute difference between the left-and right-hand terms defined as the “Distance to Criticality Coefficient” (DCC) [19]*.

*Evaluating the DCC across synchrony levels confirmed that neuronal dynamics approach a critical state at intermediate levels of synchrony in both hemispheres (Fig. 2A-B), a trend that persists even when considering large time windows (Supp. Fig. 1). As a proxy for criticality, we defined the critical region as the range in which DCC values fell below the median of their distribution. Note that at high levels of synchrony (<ρ> ≥* 0.15), a less frequently occupied state (see histogram in Fig. 1B), DCC was often undefined because the dynamics were dominated by a few large avalanches.

**FIG. 2.**
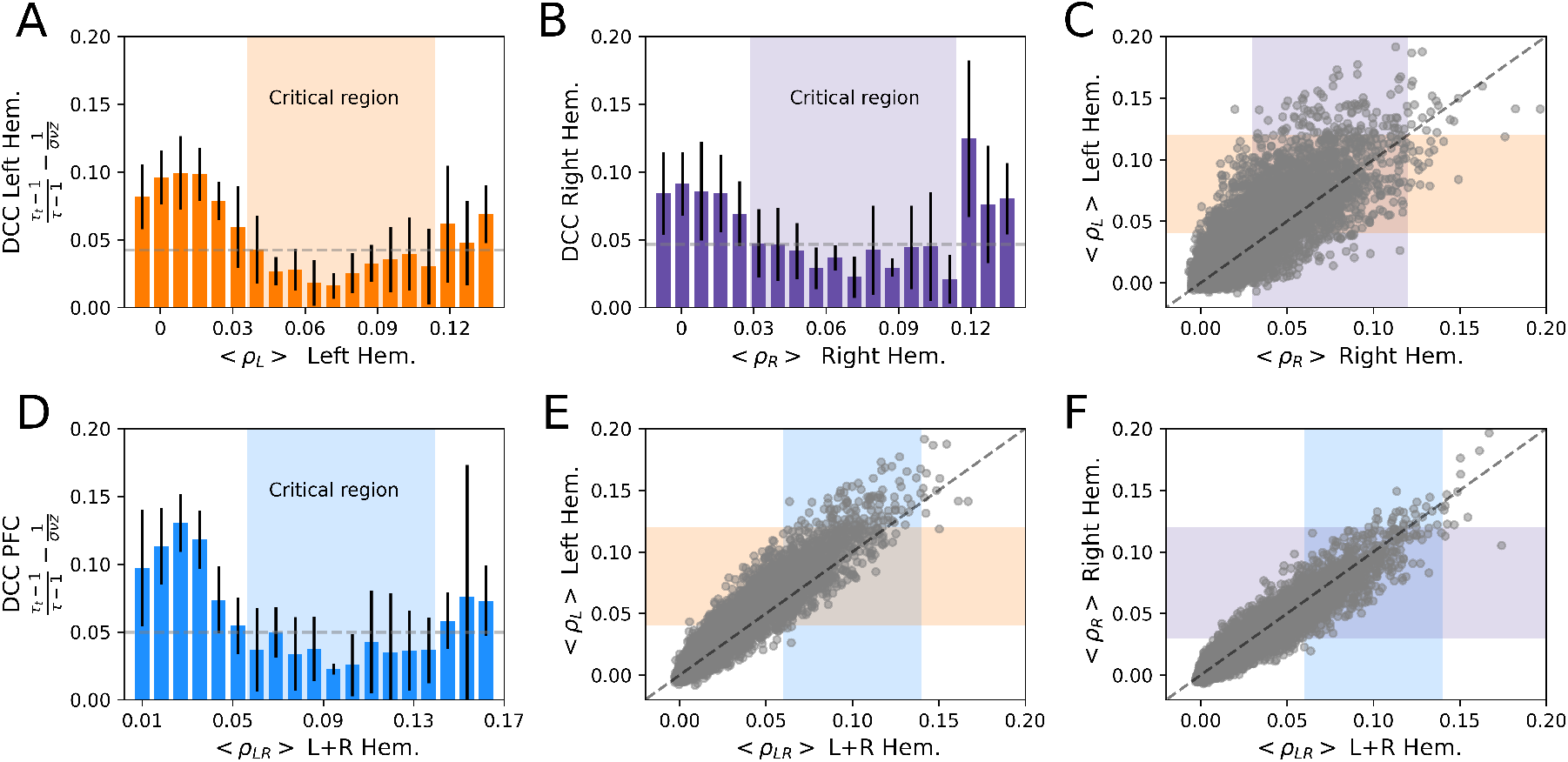
Distance to Criticality Coefficient (DCC) indicates a critical region at the edge of synchrony. (A), (B) DCC as a function of the population synchrony level, defined as the mean pairwise spiking correlation (*<ρ>*), for the left (*<ρ*_*L*_*>*; orange) and right (*<ρ*_*R*_*>*; purple) hemisphere populations, respectively. Error bars represent the standard deviation of DCC across 6 experiments (*n* = 3 subjects, each with *n*_*s*_ = 2 sessions). Shaded area marks the critical regions where DCC falls below the threshold, defined as the median of the distribution. (C) Scatter plot of *<ρ>*(*t*) for the left versus right hemispheres. Shaded areas are the same as described in (A) and (B). (D) DCC as a function of *<ρ>* for the merged left and right PFC populations (*<ρ*_*LR*_*>*). (E), (F) Scatter plot of *<ρ>*(*t*) for the left (E) and right (F) populations versus the merged population. Dashed line in (C), (E), and (F) indicates where x=y. Data in (C), (E), and (F) is from one representative rat session, the same used in Fig. 1.

Thus, our analysis showed that each PFC population approached the critical regime at intermediate levels of synchronization. We refer to this as *local criticality*, because the analysis was performed independently in the left and right populations. This finding aligns with previous theoretical work, where critical neuronal avalanches have been described at the edge of synchrony [22, 26, 27]. However, it is not yet known whether local critical fluctuations align across distant regions. Put differently, when examined within the same time window, do left and right PFC populations converge on the same near-critical state, or can they reside in distinct–and potentially opposite–regimes such as subcritical versus supercritical?

### Asymmetric Relationship Between Local and Global Criticality

To evaluate the dynamic regime between distant populations, we compared *<ρ*_*L,R*_*>*(*t*) of the left and right PFC within each time window. As illustrated in Fig. 1B, the synchronization profiles of the left and right PFC closely tracked one another [28]. In particular, we observed that intermediate levels of synchrony typically occurred simultaneously in both populations (Fig. 2C). Nonetheless, at times, we could also observe only one population in the critical state (vertical and horizontal shaded areas in Fig. 2C). These observations are consistent with evidence that, despite the structural similarities between the homotopic regions, the functional operations of the left and right PFC may diverge at certain times [29–31].

To determine whether the apparent coordination between distant populations could arise by chance, we generated 1000 surrogate datasets by shuffling *<ρ*_*R*_*>*(*t*) time series (one value per time window), thereby preserving intrapopulation correlations while disrupting interpopulation correlations. By comparing the occupancy of each combination of dynamic states (i.e., each quadrant in Fig. 2C), we found that the empirical data exhibited significantly more co-occurrence of matching states than any of the surrogates (Supp. Fig. 2). Thus, the observed interhemispheric coordination reflects genuine interactions rather than a statistical coincidence.

These observations raise the question of whether critical dynamics emerge not only locally within each hemisphere but also globally when both populations are considered as one. To address this, we repeated the same analysis, merging the left and right PFC into a single population, referred to as L+R. Using the DCC, we found that criticality was again more pronounced at intermediate levels of synchrony (Fig. 2D), consistent with the results obtained when analyzing the regions separately (Fig. 2A-B). We refer to this as *global criticality*, because the analysis was performed by merging both the left and right populations. This finding raised a further question: does global criticality require simultaneous criticality in each hemisphere, or can it emerge independently of the local states?

To address this question, we compared the synchrony of the left and right PFC populations, *<ρ*_*L,R*_*>*(*t*), with that of the merged population, *<ρ*_*LR*_*>*(*t*). We observed instances in which criticality emerged globally, i.e., in the merged population, without being present locally in either hemisphere (compare the vertical and horizontal shaded areas in Fig. 2E-F). Conversely, there were moments when one or both hemispheres exhibited critical dynamics independently, yet the merged population did not (Fig. 2E-F). These results indicate that local (within each population) and global (merged populations) signatures of criticality do not always coincide, highlighting the complex coordination of critical dynamics across scales.

### Intra- and interhemispheric Correlations Exhibit Structured Coordination

Large-scale brain studies have shown that cortical regions participate in long-range interactions across the connectome, alternating between integrated and segregated phases [28, 32–37]. Here, we hypothesized that similar long-range interactions might underlie the fluctuations in neuronal population dynamics observed in our recordings, including the fluctuations around criticality (Fig. 1 and 2). To quantify these interactions, and as a proxy of integration [38], we computed interhemispheric correlations between neurons in the two PFC regions, excluding intrahemispheric correlations. Therefore, time windows with low interhemispheric correlation were considered segregated states, whereas those with high values reflected integrated activity across populations.

We found that interhemispheric correlations followed a smooth, nontrivial gradient that depended on the intrahemispheric correlation levels within the left and right PFC (Fig. 3A, left). In other words, epochs with stronger intrahemispheric correlations were associated with higher interhemispheric correlations. To determine whether this gradient could arise trivially from increased synchronous events, we generated 1000 surrogate datasets by randomly permuting time bins within each 5-s epoch independently for the two populations. This preserved intrahemispheric correlations while disrupting interhemispheric dynamics (Fig. 3A-B). In the surrogates, both the gradient and the overall magnitude of inter-hemispheric correlations was disrupted, confirming that the empirical dependence reflects genuine long-range interactions and that interhemispheric correlation systematically depended on intrahemispheric synchrony (Fig. 3A-B).

**FIG. 3.**
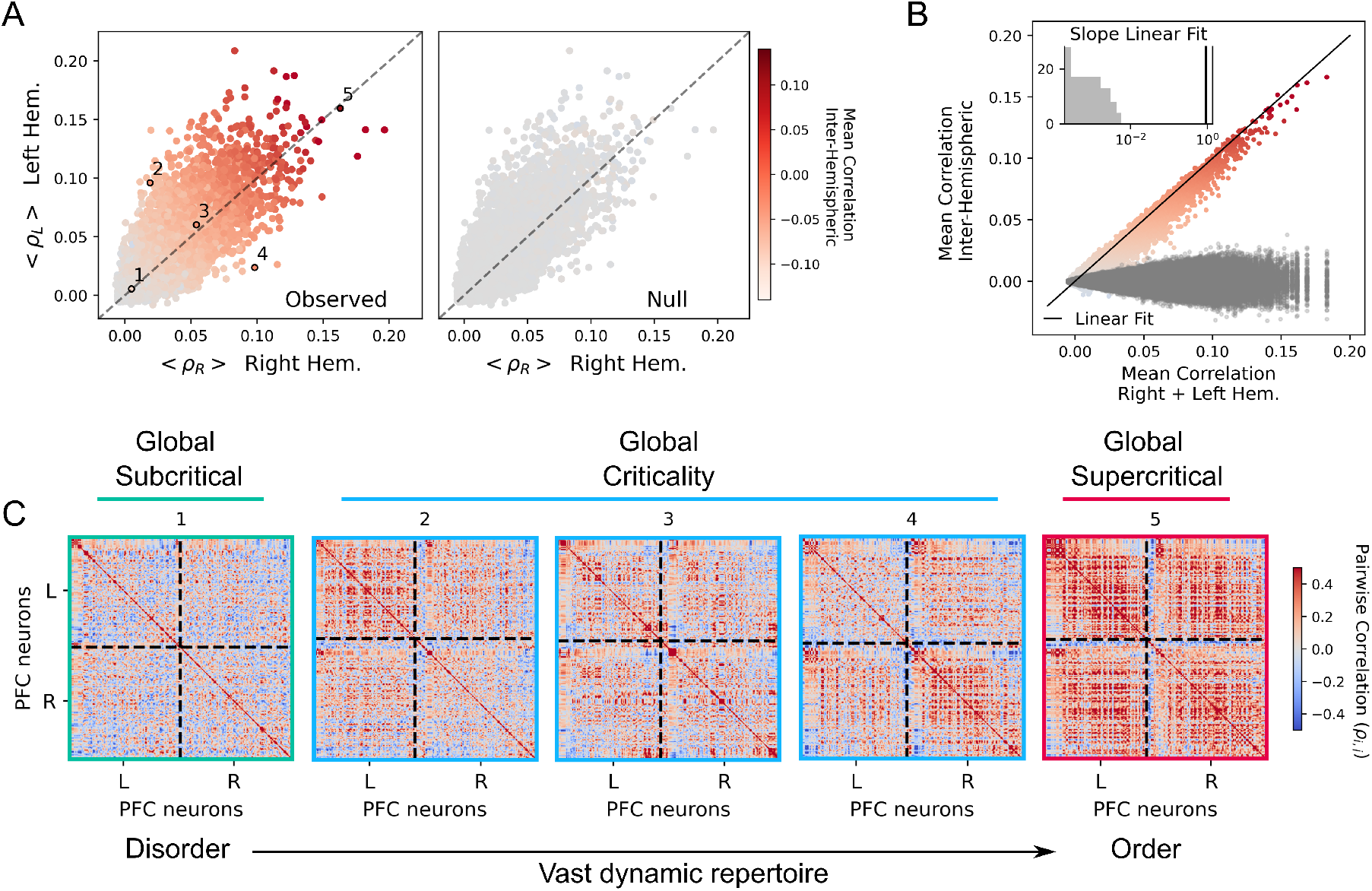
Interhemispheric correlations depends on the level of intrahemispheric correlations. (A) Scatter plots of intrahemispheric synchrony, measured as mean pairwise correlations within the left (*<ρ*_*L*_*>*) and right (*<ρ*_*R*_*>*) hemisphere populations. Each dot represents a 5-s window, color-coded by interhemispheric correlations magnitude, excluding intrahemispheric correlations. In the observed data (left), *<ρ*_*L*_*>* and *<ρ*_*R*_*>* tend to fluctuate together, with stronger interhemispheric correlations emerging when both are locally synchronized. In the null model (right), generated by shuffling time bins within each epoch independently for the two populations, this structured relationship is abolished. Numbers (1–5) correspond to representative windows shown in (C). (B) Mean interhemispheric correlations increase systematically with mean intrahemispheric correlations (average of left and right populations) in the observed data (colored points). This dependence disappears in the null model (gray points). The inset shows the distribution of slopes obtained from surrogate datasets (gray), demonstrating that the observed slope (black line) lies far outside the null distribution. (C) Representative correlation matrices from the five epochs highlighted in (A). An intrahemispheric correlation value (i.e., the *x* or *y* coordinate of a data point in panel (A) corresponds to the average value of a diagonal block, while an interhemispheric correlation value (i.e., the color scale in panel (A)) corresponds to the mean of the off-diagonal blocks. The five matrices illustrate different scenarios: scenario 1 shows low interhemispheric correlation (*<ρ>* = 0.0004) associated with low intrahemispheric correlations; Scenarios 2, 3, and 4 display moderate interhemispheric correlation (*<ρ>* = 0.045 for all three), with varying patterns of intrahemispheric correlations: scenario 2 shows higher intrahemispheric correlations in the left hemisphere, scenario 3 shows moderate intrahemispheric correlations distributed evenly across both hemispheres and scenario 4 shows higher intrahemispheric correlations in the right hemisphere; Scenario 5 represents high interhemispheric correlations (*<ρ>* = 0.139) accompanied by high intrahemispheric correlations in both hemispheres. Data shown is from one representative rat, the same used in Fig. 1.

Hence, the resulting gradient (Fig. 3A, left) indicates that local intrahemispheric synchrony shapes interhemispheric coupling. Specifically: i) when both hemispheres showed low intrahemispheric correlation, corresponding to a subcritical state, the two regions remained segregated, with minimal interhemispheric correlations (Fig. 3C, scenario 1); ii) when both hemispheres were highly synchronized, corresponding to a supercritical state, they entered an integrated state with high interhemispheric correlations (Fig. 3C, scenario 5); and iii) intermediate levels of interhemispheric correlations, reflecting a balanced state between integration and segregation, occurred when intrahemispheric correlations were intermediate in both regions (Fig. 3C, scenario 3), i.e., near a critical state, as well as when one region displayed high intrahemispheric correlations while the other remained desynchronized (Fig. 3C, scenarios 2 and 4). In particular, in scenarios 2-4, the interhemispheric correlations (i.e., the average of the off-diagonal block) were nearly the same (*<ρ>* ≃ 0.045). Therefore, these findings reveal a nontrivial aspect of cortical dynamics: neurons can exhibit significant interhemispheric correlations (global communication) without requiring high intrahemispheric synchronization (local communication), a phenomenon that, to our knowledge, has not been reported previously.

### Computational Model Near Criticality Reproduces Empirical Patterns

To test whether a system with a known critical point could account for the correlation structure observed in the data (Fig. 3A), we described each population as a network of *N* all-to-all coupled ‘active rotator’ neurons, parametrized by values for the neurons’ intrinsic drive (*ω*), noise amplitude (*σ*), and excitability (*a*) [39]. Despite its simplicity, the network of coupled active rotators reproduces three canonical regimes—subcritical (asynchronous and fluctuation-driven), critical (long-timescale, maximal susceptibility), and supercritical (oscillatory) [40] (Fig. 4A). Here, we fixed *ω* = 1 and *σ* = 0.499 as in [40], and used *a* as the control parameter. In this model, the critical point is reached at intermediate values of excitability (*a* = 1.07), where neuronal avalanches and scaling relations have been shown to emerge [40].

**FIG. 4.**
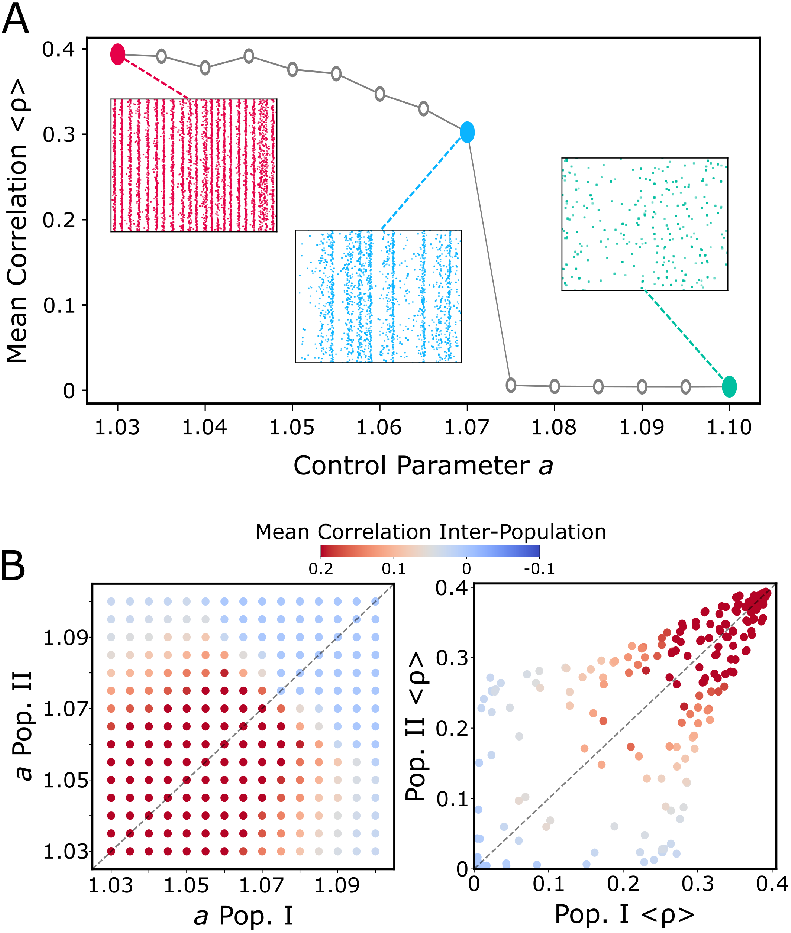
Computational model reproduces non-trivial intra- and intercorrelations. (A) Population synchrony, measured as the mean pairwise correlation *< ρ >* as a function of the control parameter *a* in the single-population model of [40]. Neurons are coupled allto-all, and the parameter *a* modulates network excitability. Example raster plots illustrating the synchronous regime (red), the intermediate regime at the model’s critical point *a* = 1.07 (blue), and the asynchronous regime (green). 250 neurons shown during 500ms. (B) (Left) Interpopulation correlations obtained by simulating two weakly coupled populations (I and II) varying the excitability parameters *a*. Color code indicates the magnitude of the mean interpopulation pairwise correlations. (B) (Right) Scatter plot of intrapopulation synchrony within population I (Pop. I *< ρ >*) and population II (Pop. II *< ρ >*). Each point corresponds to one pair (*a*_*Pop*.*I*_, *a*_*Pop*.*II*_), colorcoded by the interpopulation correlations magnitude. The dashed line marks the diagonal where *a*_*Pop*.*I*_ = *a*_*Pop*.*II*_.

We then extended the model to two weakly coupled populations (populations I and II) as an analogy to experimental left and right hemispheres populations. By varying the control parameter *a* independently in each population, we mapped the resulting interpopulation correlations (Fig. 4B left). We observe that variable levels of integration (measured as interhemispheric correlation) can be achieved by varying the decoupled local population regime around a critical state. Thus, flexible integration/segregation dynamics can be explained by variations of local critical states and longrange coupling. Notably, analyzing the simulations across this parameter space, the nontrivial intercorrelation gradient emerged (Fig. 4B right), mirroring the pattern observed in our experiments (Fig. 3A). These results demonstrated that weakly coupled networks, which hover near a critical point of an order–disorder phase transition, naturally reproduce the nontrivial relationship between local and long-range correlations observed empirically.

## Discussion

Our results demonstrate that critical dynamics in the prefrontal cortex of freely behaving rats depend on the instantaneous level of population synchrony. When examined independently, each hemisphere displayed scale-free avalanche statistics and consistent scaling relations specifically at intermediate synchrony levels (Fig. 1) [17, 25]. While high synchrony collapsed avalanche-like activity into a few large events, low synchrony produced irregular activity without signatures of criticality. These findings support the view that spontaneous cortical activity fluctuates around a critical regime, rather than *statically* operating at a critical point, and that signatures of criticality are observed at the edge of synchronization [18, 22, 26, 41].

A key finding is that local and global criticality do not necessarily align (Fig. 2). The merged left plus right populations occasionally expressed critical dynamics (*global criticality*) even when neither hemisphere was individually critical, and conversely, one or both hemispheres sometimes displayed criticality (*local criticality*) while the combined activity did not. This dissociation indicates that criticality does not scale trivially with population size [9]. Instead, it depends on both local features and global interactions, highlighting the multiscale structure of spontaneous cortical activity [42].

Long-range interactions between hemispheres further revealed a structured, nontrivial relationship between local synchrony and global coordination (Fig. 3). Interhemispheric correlations increased smoothly with intrahemispheric synchrony, and surrogate analyses confirmed that this reflected genuine functional coupling rather than trivially resulting from increased local co-activation strength. Notably, intermediate levels of interhemispheric correlations emerged not only when both hemispheres exhibited intermediate synchrony but also when one hemisphere remained desynchronized. This asymmetric configuration shows that large-scale coordination does not require matched internal states, and that the cortex can maintain cross-regional integration even when local dynamics differ.

To probe the dynamical origin of this gradient, we implemented a model with a known critical point [40]. In this model, the control parameter *a* regulates proximity to criticality, and variations in *a* can be interpreted as reflecting spontaneous neuromodulation effects as well as slow changes in arousal or behavioral state. When two populations were weakly coupled through long-range interactions, modulating *a* was sufficient to reproduce the nontrivial gradient observed experimentally, suggesting that coordinated fluctuations in synchrony can emerge naturally in systems whose effective dynamics are shaped by local, state-dependent proximity to criticality [13, 20, 43].

Together, our results support a multiscale view of criticality in which local synchrony shapes, but does not fully determine, global network behavior. In this view, criticality provides a regime that maximizes flexibility: it enables local populations to explore diverse spatiotemporal patterns while allowing dynamic coupling across regions (Fig. 3C), consistent with accounts proposing that networks near a continuous phase transition balance segregation and integration [24, 38, 44, 45].

More broadly, our findings extend neuronal signatures of criticality beyond isolated cortical populations and emphasize it as a potential organizing principle for distributed cortical interactions. However, our study also has limitations. For instance, we focused on homotopic PFC regions, which, despite at times exhibiting distinct temporal dynamics, often fluctuated in unison, likely reflecting their shared structural connectivity and computational roles [30, 36, 46, 47]. As large-scale electrophysiological and imaging approaches continue to expand [48, 49], it will be important to assess whether these principles generalize to more heterogeneous networks. Prior work suggests that distance from criticality may vary hierarchically across brain areas [10, 50], raising the possibility that different regions contribute differently to multiscale coordination. However, it is still unclear whether these variations reflect local properties or emerge from dynamic interactions across the connectome. Large-scale computational models have begun to address these questions, linking local to global brain dynamics. In particular, the accompanying paper [51] introduces a multiscale model displaying the coemergence of local and global near-critical dynamics, and provides the first in silico evidence that local distance from criticality is shaped both by the local properties and by the long-range interactions over the connectome. Altogether, exploring how critical dynamics develop on multiple scales may reveal their role in supporting flexible computations and how they relate to neurological and psychiatric (dys)function [44, 52–54

## Methods

### Data acquisition and processing

Recordings from three male Long-Evans rats (350–400 g at the time of surgery) were used for this study. All data in this study come from a previously published study [55]. Full details about animal care, surgery, data acquisition, and processing can be found in [55, 56]. In compliance with European Directives 86*/*609*/*CEE and 2010*/*63*/*UE, all experiments were evaluated by an ethics committee (CEEA #59) and approved by the French Ministry of Research (authorization #8229).

Briefly, the animals were housed individually in monitored conditions (21^°^C and 45% humidity) and maintained on a 12h light – 12h dark cycle. To avoid obesity, food was restricted to 13–16 g of rat chow per day, while water was available ad libitum. To habituate the rats to human manipulation, they were handled each workday. 16-24 bundles of twisted-wire electrodes were implanted in the PFC 2.5 to 4.8 mm anterior and ±0.3 to 0.6 mm lateral from bregma with reference and ground screws in the cerebellum. Brain activity was recorded with a 256-channel digital data acquisition system (KJE-1001, Amplipex, Szeged, Hungary).

The animals underwent a multi-day habituation, fear conditioning, and recall test protocol as described in [56], and the two sessions per animal used here correspond to the habituation and conditioning phases. The signals were continuously acquired over ∼8*h* per session with four 64-channel headstages (Amplipex HS2) and sampled in wideband at 20 kHz. An inertial measurement unit sampled the 3D angular velocity and linear acceleration of the head at 300 Hz. Off-line spike sorting was performed using Kilosort 1.0 [57]. The resulting clusters were visually inspected, using Klusters [58], to reject noise and to merge units erroneously considered as different units. The total number of single units registered in the right (left) hemisphere of the three animals was: 127 (133), 78 (129), and 81 (72).

### Time Window Segmentation

Continuous spiking data from each PFC population were divided into non-overlapping 5-second segments to capture temporal fluctuations in population activity, unless otherwise stated. Each 5-second window was treated independently for subsequent analyses, including the computation of mean pairwise spiking correlations and neuronal avalanches. This segmentation allowed us to examine dynamic changes in synchrony and criticality over time while maintaining sufficient temporal resolution to detect avalanches.

### Pairwise Spike Correlations

To compute pairwise spiketrain correlations *ρ*, spike trains *s*(*t*) were discretized using fixed 20-ms time bins and then convolved with a kernel *k*:

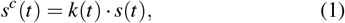

where *k* is a Mexican hat kernel defined as the difference between two zero-mean Gaussian distributions with standard deviations of 100 ms and 400 ms, respectively [18, 59]. From the convolved spike trains, *s*^*c*^(*t*), we computed the pairwise correlation between units *i* and *j* as:

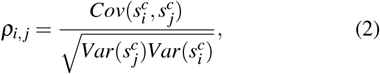

where *Cov* denotes covariance and *Var* denotes variance. The mean pairwise correlation, *<ρ>*, was used as a measure of population-level synchrony.

### Neuronal Avalanches Analysis

Spiking activity from simultaneously recorded neurons was binned in time using a bin width Δ*t* defined as the mean interspike interval of the population, following the original approach by Beggs and Plenz [5]. Each 5-s recording segment was analyzed separately to capture fluctuations in population dynamics over time. An avalanche was defined as a sequence of consecutive time bins containing at least one spike, bounded before and after by empty bins. The size of an avalanche, *S*, was calculated as the total number of spikes occurring across all occupied bins in the event. The duration, *T*, was defined as the number of consecutive occupied bins. Avalanches were then grouped according to synchrony level using 30 non-overlapping bins spanning the full observed range of *<ρ>*(*t*).

To fit the power-law distributions of avalanche sizes *P*(*S*) ∼ *S* ^−*τ*^ and durations 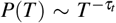, and to obtain their corresponding critical exponents, a maximum likelihood estimator was used [15, 18]. As a stringent test of critical dynamics, the scaling relation between avalanche size and duration, ⟨*S*⟩(*T*) ∼ *T* ^1*/σνz*^, was compared with the expected relation 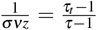 [17]. The absolute difference between these two quantities was defined as the “Distance to Criticality Coefficient” (DCC) [19]. DCC was computed for each segment and assessed across the observed range of population synchrony to identify intervals in which the system approached the critical regime, defined as periods when the DCC fell below the median of its distribution. Despite the heuristic nature of this threshold, our results clearly reveal a U-shaped relationship between DCC and synchrony levels, indicating a pronounced minimum at intermediate levels of synchrony (Fig. 2).

### Null Models

To assess whether coordination between populations could arise by chance, we generated 1000 null models by shuffling the pairwise spike train correlations time series of the right hemisphere (*<ρ*_*R*_*>*(*t*)). This procedure preserved the intrahemispheric correlations within each segment while disrupting temporal alignment between the left and right populations, thereby eliminating interhemispheric synchrony. The probability of occupancy for each possible combination of dynamic states was then computed for both the empirical data *<ρ*_*L*_*>* versus *<ρ*_*R*_*>* and versus the shuffled time series *<ρ*_*R,shu f f led*_*>*.

To assess statistical significance, we compared each empirical (observed) value to its corresponding null distribution. For each case, we first computed the mean of the null distribution. We then calculated the absolute deviation of the observed value from this null mean. An empirical two-tailed p-value was obtained as the proportion of null samples whose absolute deviation from the null mean was greater than or equal to that of the observed value. Statistical significance was assessed at the *α* = 0.05 level.

To construct the null model used in Fig. 3, we first represented the spiking activity of each population as a matrix of neurons (rows) by time (columns). We then generated 1000 null models by randomly permuting the time bins (i.e., columns) of this matrix, independently for the two hemispheres. Because the permutation was applied to entire columns, the instantaneous correlation structure among neurons within a hemisphere was preserved, and temporal alignment across hemispheres was disrupted. This procedure, therefore, disrupted interhemispheric correlations while preserving intrahemispheric correlations. Slopes shown in Fig. 3B were obtained using linear regression implemented in the SciPy Python package.

### Computational Model

We modeled each region *n* as a network of *K* = 250 all-to-all interconnected neurons, represented as active rotators governed by the Shinomoto-Kuramoto model, as described in [39, 40]:

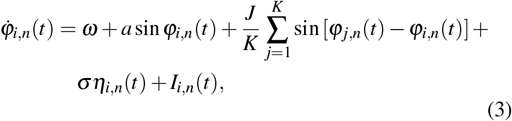

where *ϕ*_*i,n*_ is the phase of the *i*-th neuron in region *n* ∈ {0, 1}, *ω* = 1 is the intrinsic frequency, and *J* = 1 is the within-region coupling strength. *a* represents the excitability of the system and serves as a control parameter for its phase transition (see Fig. 2 in [40]). Additionally, each neuron receives an independent Gaussian white noise *η*_*i,n*_(*t*) of amplitude *σ* = 0.499. The connection across populations, i.e., long-range coupling, enters as an additive term:

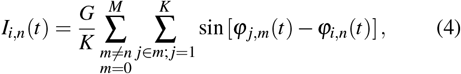

where the first summation indexes the population (*M* = 1), and the second summation accounts for all neuron pairs contributing to the long-range coupling. *G* = 0.1 is the *global coupling* parameter.

For *G* = 0, Eq. (3) describes a disconnected population. In this case, for appropriated values of *a* and *σ*, the model exhibits a hybrid-type (HT) synchronization transition [40, 60], in which asynchronous states and incipient oscillations coexist, marking a critical transition between asynchronous and synchronous regimes. At the critical point of this transition, the sizes and durations of neuronal avalanches follow power-law distributions, and the scaling relations predicted for criticality are satisfied [40]. For a detailed phase diagram of this model, refer to [40]. In this model, neuronal activity was defined as *y*_*i,n*_(*t*) = sin *ϕ*_*i,n*_(*t*) and a spike was defined as every time *y*_*i,n*_(*t*) crossed a threshold *y*_*th*_ = 0.6 from below [40]. The resulting spike train was analyzed in the same way as the empirical spiking data to compute pairwise correlations. Simulation time was rescaled by a factor of 1*/*1000 to obtain realistic unit firing rates, i.e., 1s of simulation was interpreted as 1ms, without loss of generality, since the model is phenomenological and the units are arbitrary. Simulations were performed in custom Python scripts using Brian2 software [61].

## Data, Materials, and Software Availability

The code supporting the findings of this study will be made publicly available upon acceptance of the manuscript. Data supporting the findings of this study will be made available from the corresponding author upon reasonable request.

## Acknowledgments

GR acknowledges support from the Marie Skłódowska–Curie Postdoctoral Fellowship (Project CAERUS) under the European Union’s Horizon Europe research and innovation programme (Grant No. 101199894). LDP acknowledges support from the European Union (ERC, NEMESIS, project number 101071900), AGAUR co-funded by the Departament de Recerca i Universitats de la Generalitat de Catalunya (AGAUR 2021−SGR−01165), and the Brazilian agency CNPq (Grant No. 444500*/*2024 − 3).

## Author contributions

L.D.P. and G.R. designed research; G.R. and P.B. performed simulations; M.N.P. performed experiments and preprocessed experimental data; L.D.P., P.B., and G.R. analyzed data; L.D.P. and G.R. organized results; L.D.P. wrote the first draft; all authors discussed the results and wrote the paper.

---

**Supplementary Figure 1.**
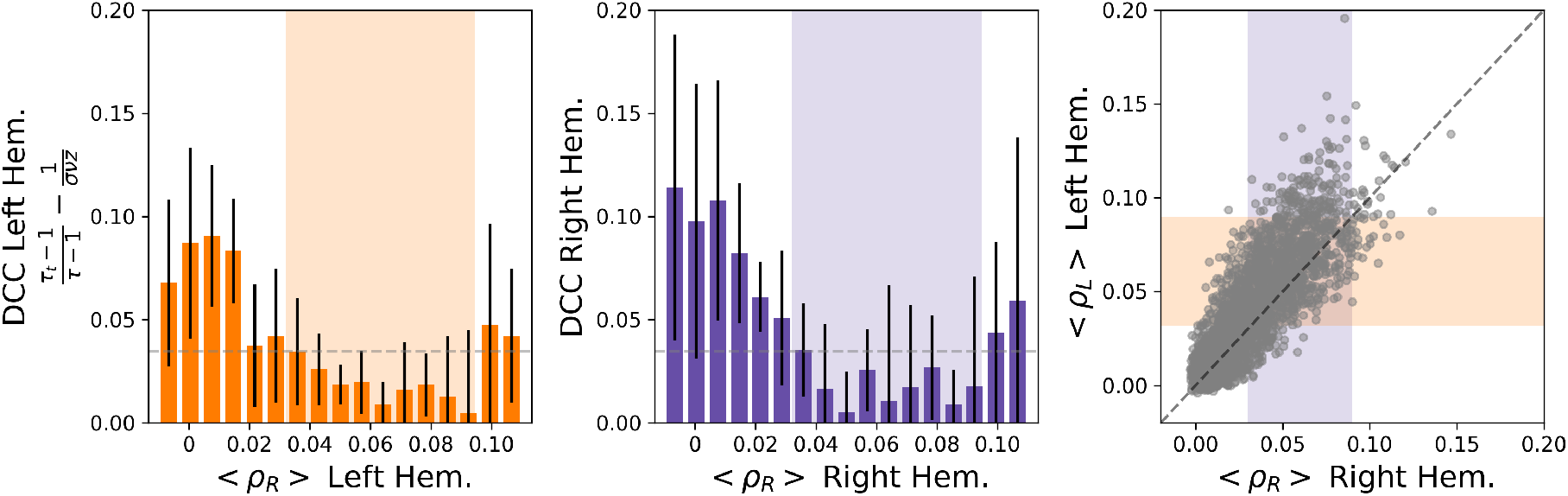
(Left, Center) Distance to Criticality Coefficient (DCC) as a function of the population synchrony level, defined as the mean pairwise spiking correlation (*<ρ>*), for the left (*<ρ*_*L*_*>*; orange) and right (*<ρ*_*R*_*>*; purple) hemisphere populations, respectively, *<ρ>* was computed over 10-seconds non-overlapping time windows (Δ*T* = 10s). Shaded area marks the critical regions where DCC falls below the threshold, defined as the median of the distribution. (Right) Scatter plot of *<ρ>*(*t*) for the left versus right hemispheres.

**Supplementary Figure 2.**
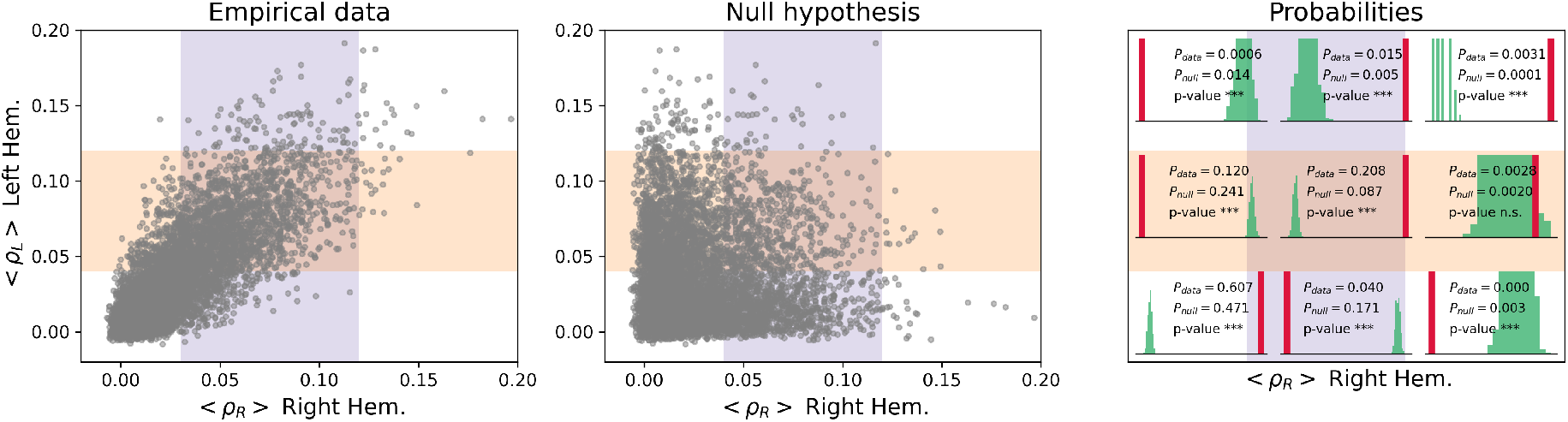
(Left) Scatter plot of *<ρ>*(*t*) for the left versus right hemispheres. Shaded areas indicate the critical region for each population: orange for the left hemisphere and purple for the right hemisphere. Same data as in Fig. 2A-C. (Middle) 1000 null models were created by shuffling *<ρ*_*R*_*>*, preserving intrahemispheric correlations but disrupting interhemispheric correlations. (Right) Occupancy probability for the empirical data (*P*_*data*_) versus the null hypothesis (*P*_*null*_). Statistical significance was defined through a permutation-based p-value with threshold *α <* 0.05.

## Notes

### Competing Interest Statement

The authors have declared no competing interest.

## References

[1] Tononi G, Sporns O, Edelman GM (1994) A measure for brain complexity: relating functional segregation and integration in the nervous system. Proceedings of the National Academy of Sciences 91(11):5033–5037.

[2] Cohen JR, D’Esposito M (2016) The segregation and integration of distinct brain networks and their relationship to cognition. Journal of Neuroscience 36(48):12083–12094.

[3] Rosen MC, Freedman DJ (2025) How distributed is the brain-wide network that is recruited for cognition? Nature Reviews Neuroscience pp. 1–13.

[4] Beggs JM (2008) The criticality hypothesis: how local cortical networks might optimize information processing. Philosophical Transactions of the Royal Society A: Mathematical, Physical and Engineering Sciences 366(1864):329–343.

[5] Beggs JM, Plenz D (2003) Neuronal avalanches in neocortical circuits. Journal of neuroscience 23(35):11167–11177.

[6] Palva JM, Palva S (2014) The correlation of the neuronal long-range temporal correlations, avalanche dynamics with the behavioral scaling laws and interindividual variability. Criticality in Neural Systems pp. 105–126.

[7] Shew WL, Plenz D (2013) The functional benefits of criticality in the cortex. The neuroscientist 19(1):88–100.

[8] Hengen KB, Shew WL (2025) Is criticality a unified setpoint of brain function? Neuron 113(16):2582–2598.

[9] Jones SA, Barfield JH, Norman VK, Shew WL (2023) Scale-free behavioral dynamics directly linked with scale-free cortical dynamics. Elife 12:e79950.

[10] Harris B, Gollo LL, Fulcher BD (2024) Tracking the distance to criticality in systems with unknown noise. Physical Review X 14(3):031021.

[11] Fontenele AJ, Sooter JS, Norman VK, Gautam SH, Shew WL (2024) Low-dimensional criticality embedded in high-dimensional awake brain dynamics. Science Advances 10(17):eadj9303.

[12] Dahmen D, Grün S, Diesmann M, Helias M (2019) Second type of criticality in the brain uncovers rich multiple-neuron dynamics. Proceedings of the National Academy of Sciences 116(26):13051–13060.

[13] Sooter JS, et al. (2025) Defining and measuring proximity to criticality. bioRxiv pp. 2025–08.

[14] Hahn G, et al. (2017) Spontaneous cortical activity is transiently poised close to criticality. PLoS computational biology 13(5):e1005543.

[15] Alstott J, Bullmore E, Plenz D (2014) powerlaw: a python package for analysis of heavy-tailed distributions. PloS one 9(1):e85777.

[16] Munoz MA, Dickman R, Vespignani A, Zapperi S (1999) Avalanche and spreading exponents in systems with absorbing states. Physical Review E 59(5):6175.

[17] Friedman N, et al. (2012) Universal critical dynamics in high resolution neuronal avalanche data. Physical review letters 108(20):208102.

[18] Fontenele AJ, et al. (2019) Criticality between cortical states. Physical review letters 122(20):208101.

[19] Ma Z, Turrigiano GG, Wessel R, Hengen KB (2019) Cortical circuit dynamics are homeostatically tuned to criticality in vivo. Neuron 104(4):655–664.

[20] Xu Y, Schneider A, Wessel R, Hengen KB (2024) Sleep restores an optimal computational regime in cortical networks. Nature neuroscience 27(2):328–338.

[21] O’Byrne J, Jerbi K (2022) How critical is brain criticality? Trends in Neurosciences 45(11):820–837.

[22] Di Santo S, Villegas P, Burioni R, Muñoz MA (2018) Landau– ginzburg theory of cortex dynamics: Scale-free avalanches emerge at the edge of synchronization. Proceedings of the National Academy of Sciences 115(7):E1356–E1365.

[23] Yu S, et al. (2017) Maintained avalanche dynamics during task-induced changes of neuronal activity in nonhuman primates. elife 6:e27119.

[24] Munoz MA (2018) Colloquium: Criticality and dynamical scaling in living systems. Reviews of Modern Physics 90(3):031001.

[25] Sethna JP, Dahmen KA, Myers CR (2001) Crackling noise. Nature 410(6825):242–250.

[26] Poil SS, Hardstone R, Mansvelder HD, Linkenkaer-Hansen K (2012) Critical-state dynamics of avalanches and oscillations jointly emerge from balanced excitation/inhibition in neuronal networks. Journal of Neuroscience 32(29):9817–9823.

[27] Dalla Porta L, Copelli M (2019) Modeling neuronal avalanches and long-range temporal correlations at the emergence of collective oscillations: Continuously varying exponents mimic m/eeg results. PLoS computational biology 15(4):e1006924.

[28] Nowak L, Munk M, Nelson J, James A, Bullier J (1995) Structural basis of cortical synchronization. i. three types of interhemispheric coupling. Journal of Neurophysiology 74(6):2379–2400.

[29] Yin X, Wang Y, Li J, Guo ZV (2022) Lateralization of short-term memory in the frontal cortex. Cell Reports 40(7).

[30] Brincat SL, et al. (2021) Interhemispheric transfer of working memories. Neuron 109(6):1055–1066.

[31] Ghosh M, et al. (2022) Running speed and rem sleep control two distinct modes of rapid interhemispheric communication. Cell reports 40(1).

[32] Sporns O (2011) The human connectome: a complex network. Annals of the new York Academy of Sciences 1224(1):109–125.

[33] Betzel RF, Bassett DS (2018) Specificity and robustness of long-distance connections in weighted, interareal connectomes. Proceedings of the National Academy of Sciences 115(21):E4880–E4889.

[34] Sauerbrei BA, et al. (2020) Cortical pattern generation during dexterous movement is input-driven. Nature 577(7790):386–391.

[35] Machado TA, Kauvar IV, Deisseroth K (2022) Multiregion neuronal activity: the forest and the trees. Nature Reviews Neuroscience 23(11):683–704.

[36] Engel AK, König P, Kreiter AK, Singer W (1991) Interhemispheric synchronization of oscillatory neuronal responses in cat visual cortex. Science 252(5009):1177–1179.

[37] Sorrentino P, et al. (2021) The structural connectome constrains fast brain dynamics. Elife 10:e67400.

[38] Deco G, Tononi G, Boly M, Kringelbach ML (2015) Rethinking segregation and integration: contributions of whole-brain modelling. Nature Reviews Neuroscience 16(7):430–439.

[39] Shinomoto S, Kuramoto Y (1986) Phase transitions in active rotator systems. Progress of Theoretical Physics 75(5):1105–1110.

[40] Buendía V, Villegas P, Burioni R, Muñoz MA (2021) Hybrid-type synchronization transitions: Where incipient oscillations, scale-free avalanches, and bistability live together. Physical Review Research 3(2):023224.

[41] Toker D, et al. (2022) Consciousness is supported by near-critical slow cortical electrodynamics. Proceedings of the National Academy of Sciences 119(7):e2024455119.

[42] Engel TA, Steinmetz NA (2019) New perspectives on dimensionality and variability from large-scale cortical dynamics. Current opinion in neurobiology 58:181–190.

[43] Castro DM, et al. (2024) In and out of criticality? state-dependent scaling in the rat visual cortex. PRX Life 2(2):023008.

[44] Cocchi L, Gollo LL, Zalesky A, Breakspear M (2017) Criticality in the brain: A synthesis of neurobiology, models and cognition. Progress in neurobiology 158:132–152.

[45] Sporns O (2013) Network attributes for segregation and integration in the human brain. Current opinion in neurobiology 23(2):162–171.

[46] Bhattacharya S, Brincat SL, Lundqvist M, Miller EK (2022) Traveling waves in the prefrontal cortex during working memory. PLoS computational biology 18(1):e1009827.

[47] Oran Y, Katz Y, Sokoletsky M, Malina KCK, Lampl I (2021) Reduction of corpus callosum activity during whisking leads to interhemispheric decorrelation. Nature communications 12(1):4095.

[48] Stringer C, et al. (2019) Spontaneous behaviors drive multidimensional, brainwide activity. Science 364(6437):eaav7893.

[49] Findling C, et al. (2025) Brain-wide representations of prior information in mouse decision-making. Nature 645(8079):192–200.

[50] Morales GB, Di Santo S, Muñoz MA (2023) Quasiuniversal scaling in mouse-brain neuronal activity stems from edge-of-instability critical dynamics. Proceedings of the National Academy of Sciences 120(9):e2208998120.

[51] Rabuffo G, et al. (2026) The connectome modulates critical brain dynamics across local and global scales. BioRxiv.

[52] Zimmern V (2020) Why brain criticality is clinically relevant: a scoping review. Frontiers in neural circuits 14:54.

[53] Müller PM, Miron G, Holtkamp M, Meisel C (2025) Critical dynamics predicts cognitive performance and provides a common framework for heterogeneous mechanisms impacting cognition. Proceedings of the National Academy of Sciences 122(14):e2417117122.

[54] Xin Y, Cui Y, Yu S, Liu N (2025) Genetic contributions to brain criticality and its relationship with human cognitive functions. Proceedings of the National Academy of Sciences 122(26):e2417010122.

[55] Pompili, Marco N, et al. (2025) Adaptive communication between cell assemblies and “reader” neurons shapes flexible 10 brain dynamics. PLOS Biology.

[56] Pompili, Marco N, Hamou N, Wiener, Sidney I (2024) Integration of fear learning and fear expression across the dorsoventral axis of the hippocampus. bioRxiv.

[57] Pachitariu M, Steinmetz NA, Kadir SN, Carandini M, Harris KD (2016) Fast and accurate spike sorting of high-channel count probes with kilosort in Advances in Neural Information Processing Systems, eds. Lee D, Sugiyama M, Luxburg U, Guyon I, Garnett R. (Curran Associates, Inc.), Vol. 29.

[58] Hazan L, Zugaro M, Buzsáki G (2006) Klusters, neuroscope, ndmanager: a free software suite for neurophysiological data processing and visualization. Journal of neuroscience methods 155(2):207–216.

[59] Renart A, et al. (2010) The asynchronous state in cortical circuits. science 327(5965):587–590.

[60] Klinshov V, Franović I (2019) Two scenarios for the onset and suppression of collective oscillations in heterogeneous populations of active rotators. Physical Review E 100(6):062211.

[61] Stimberg M, Brette R, Goodman DF (2019) Brian 2, an intuitive and efficient neural simulator. elife 8:e47314.

